# *In situ* solid-liquid extraction enhances recovery of taxadiene from engineered *S. cerevisiae* cell factories

**DOI:** 10.1101/2021.07.12.452013

**Authors:** Jorge H. Santoyo-Garcia, Laura E. Walls, Behnaz Nowrouzi, Marisol Ochoa-Villareal, Gary J. Loake, Simone Dimartino, Leonardo Rios-Solis

## Abstract

A novel *in situ* solid phase adsorption strategy was investigated for enhanced recovery of taxadiene, a precursor to the blockbuster anticancer drug, paclitaxel, from engineered *Saccharomyces cerevisiae*. A synthetic adsorbent resin (HP-20) was employed to capture taxadiene across a range of cultivation scales. Cultivations from 12 % (w/v) resin concentration resulted in bead fragmentation which were found to be detrimental to cellular growth. After cultivation, the use of acetone for desorption captured intracellular and secreted taxadiene, achieving an integration of the bioprocess. Implementation of the proposed method at microscale (2 mL) and benchtop bioreactor scale (250 mL) resulted in 1.9-fold and 1.4-fold increments in taxadiene titer, respectively, compared to the extraction method using a dodecane overlay. Taxadiene was found to be distributed between resin beads and biomass in a ratio of 50 %. Finally, a maximum taxadiene titer of 76 ± 19 mg/L was achieved in the benchtop bioreactor cultivations.

## 1. Introduction

Downstream processing for the recovery and purification of valuable natural products (NPs) typically accounts for 60-80% of the total operational costs at industrial scale [1]. Such processing steps are typically energy intensive and require vast quantities of highly toxic solvents, resulting in major economic and environmental sustainability challenges. NPs have extensive applications across the global food, cosmetics and pharmaceutical industries [2,3]. As a result of such far-reaching potential benefits, NPs consumption has tripled since the 1990s [4].

Over 60% of all known NPs are terpenoids, a large chemically diverse group of compounds with important roles in both primary and secondary metabolism in their natural hosts [5,6]. Some terpenoids in particular are drug candidates for the treatment of human disease; examples include the sesquiterpenoid anti-malarial drug, artemisinin and diterpenoid chemotherapy drug, paclitaxel [7,8]. Most terpenoids are hydrophobic in nature and often become toxic to heterologous hosts at industrially feasible concentrations [9]. To minimize the detrimental effects of terpenoids on the host, *in situ* product recovery methods have become common practice at laboratory scale [10,11]. Through *in situ* extraction, terpenoids can be effectively removed from the cultivation medium as they are produced, thereby minimizing the accumulation, product loss and cell death [12].

### 1.1 *In situ* solid phase adsorption of natural products

*In situ* solid phase adsorption (SPA) is an alternative method, which could potentially alleviate the aforementioned bottlenecks associated with LLE. During *in situ* SPA, an inert adsorbent resin is added to the cultivation medium and the NP adsorbs to its surface as it is secreted by the host [13]. This dramatically reduces product loss due to air stripping or co-evaporation with the organic overlay [14].

Through *in situ* SPA, feedback inhibition and cytotoxic effects can be minimized thereby enhancing productivity [15,16]. Adsorbent resins have been successfully employed in this context to extract a wide range of NPs from a number of heterologous hosts including *Escherichia coli*, *Aspergillus fumigatus* and *Streptomyces hygroscopicus* via *in situ* or post-cultivation adsorption [15–18]. Different SPA processes have been developed, one example is the extraction of chaetominine, an alkaloid compound with anti-cancer activity, from *Aspergillus fumigatus*, where out of eight resins tested, XAD-16 and HP-20 demonstrated the best extraction performance [17]. This superior performance was attributed to greater adsorption capacity resulting from their larger surface area. A number of polymeric resins have also been investigated recently for the recovery of the sesquiterpene, (+)-zizaene, from engineered *E. coli* cultivations. Of those tested, the highest (+)-zizaene titer was achieved using HP-20 in the *E. coli* cultivations, where the resin structure was a key element for higher hydrophobic adsorption [19].

### 1.2 Research scope: *In situ* solid phase adsorption for taxadiene biosynthesis and recovery

The biosynthesis and recovery of the highly effective chemotherapy drug, paclitaxel (commercially known as Taxol ^®^), is a major ongoing research effort [20,21]. Despite decades of study, the complex paclitaxel biosynthetic pathway is yet to be fully elucidated. A robust precursor purification method therefore has the potential to expedite cell factory development and hence progress towards a more sustainable source of the drug [7,22]. Heterologous biosynthesis of the first committed precursor in the paclitaxel pathway, taxadiene, has been successfully achieved in both *E. coli* [23,24] and *S. cerevisiae* [25,26].

The use of a dodecane overlay has proven to be an effective method for taxadiene recovery in the highest-yielding *E. coli* (Ajikumar et al., 2010; Biggs et al., 2016) and *S. cerevisiae* at laboratory scale cultivations [21,25]. However, research into more scalable recovery and purification methods for this critical precursor is scarce, despite taxadiene itself being considered a valuable NP [22]. As taxadiene is a hydrophobic molecule by nature, a non-polar resin is appropriate for its extraction. Of the wide range of available non-polar resins, HP-20 beads (Diaion^®^) were deemed most suitable for taxadiene recovery due to their very large surface area (850-1000 m^2^/g) and high adsorption capacity of NPs with similar chemical structure (chaetominine) [17].

The metabolic pathway for heterologous production and subsequent adsorption of taxadiene onto the HP-20 resin beads is summarized in Figure 1. Although HP-20 resin beads have been employed for the adsorption and subsequent recovery of a variety of NPs, their application for the extraction of microbially synthesised taxadiene is novel.

**Figure 1.**
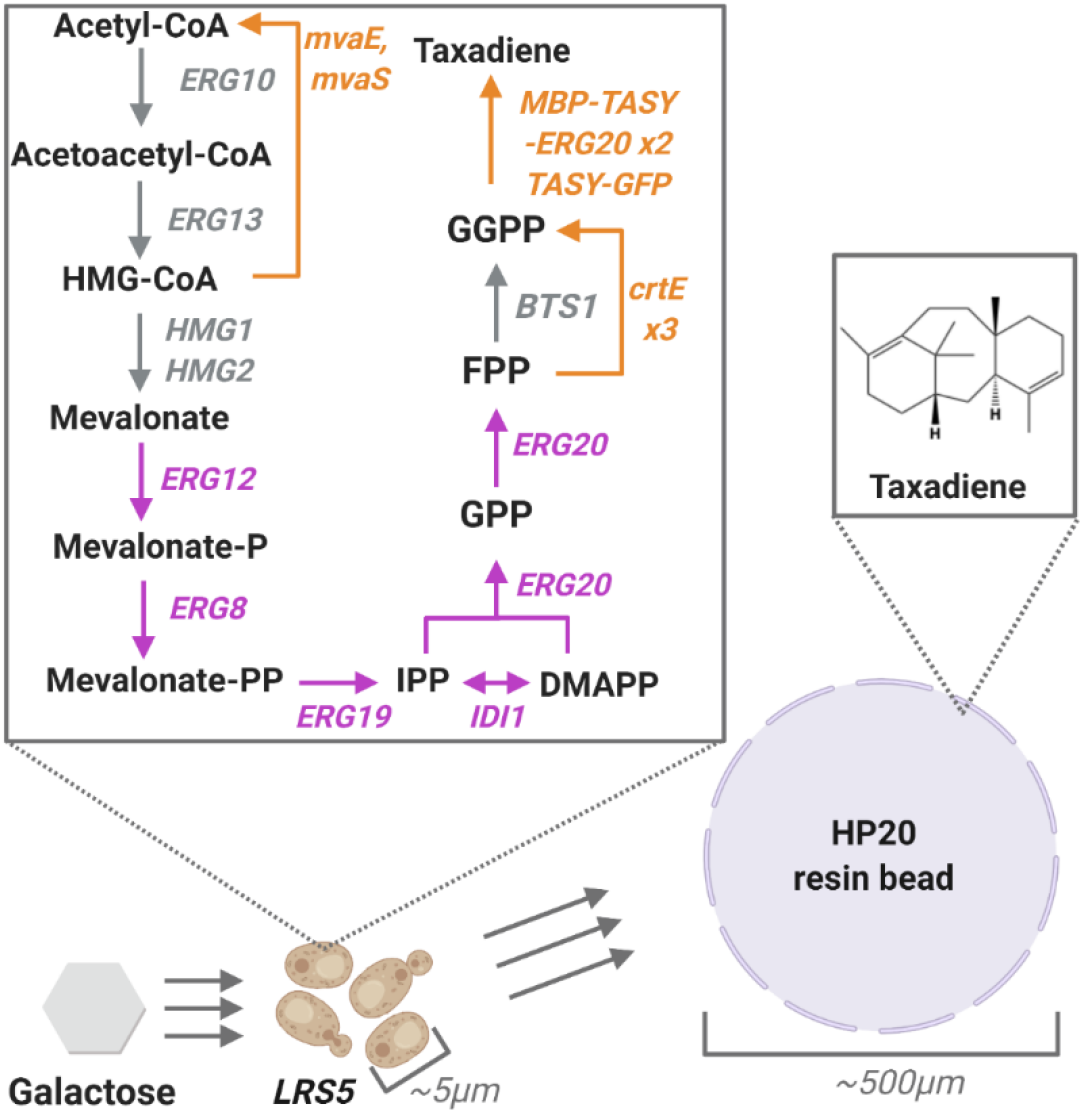
Heterologous taxadiene biosynthetic pathway in *S. cerevisiae* engineered strain (*LRS5*) and concurrent *in situ* adsorption of the secreted taxadiene to the surface of the HP-20 resin beads. Galactose is metabolized via glycolysis yielding acetyl-CoA, which is then converted into the universal diterpenoid precursor, geranylgeranyl diphosphate (GGPP), via the highlighted mevalonate pathway. Finally GGPP undergoes cyclization by taxadiene synthase (TASY) yielding taxadiene. Native overexpressed genes are highlighted in purple, whilst heterologous genes are shown in orange.

In this study, a HP-20 resin bead mediated SPA method for the recovery of taxadiene produced by an engineered *S. cerevisiae* strain, *LRS5* [25], was developed and optimized. The effect of key factors such as resin concentration and extraction solvent on product recovery were investigated in detail across a range of bioreactor scales. The traditional liquid-liquid extraction approach was used as a control to assess performance. The system was tested across a range of cultivation scales from 2 mL high throughput cultivations previously shown to be predictive of larger scale operations [11], to controlled benchtop bioreactors (250 mL) to investigate scalability and performance under more industrially relevant conditions. Finally, the bioprocess was intensified by provoking cell lysis using cell-wall disruptive organic solvents in the final extraction step.

## 2. Materials and Methods

### 2.1 Yeast strains and media

The *S. cerevisiae* strain employed in this study was *LRS5 {mGTy116; ARS1014::GAL1p-TASY-GFP; ARS1622b::GAL1p-MBP-TASY-ERG20; ARS1114a::TDH3p-MBP-TASY-ERG20}* as described in detail in [25] was originally derived from laboratory strain CEN.PK2-1C (EUROSCARF, Germany). All reagents, including extraction solvents were sourced from Fisher Scientific UK at the highest available purity unless otherwise stated. The media used for the yeast cultivations was yeast extract peptone (YP, yeast extract 1 % (w/v), peptone 2 % (w/v)) supplemented with 2 % (w/v) galactose (YPG) or 2 % (w/v) glucose (YPD) unless otherwise stated.

### 2.2 Microscale cultivation using the solid phase adsorption (SPA) method

High-throughput microscale screening was performed using 24-well 10 ML deep well plates (Axygen, USA) with a working volume of 2 mL. Inoculum cultures were prepared by transferring a single colony of *LRS5* to 5 mL of YPD media and incubating at 30 °C and 250 rpm overnight. An aliquot of this culture was then diluted with YPG media to give a 2 mL culture with an initial OD_600_ = 1. Autoclaved HP-20 adsorbent resin beads were added to each of the cultivations at concentrations between 3 to 12 % (w/v) as indicated in the loading plan (see supplementary material). Prior to use, the beads were autoclaved, rinsed with 100% ethanol and washed with sterile deionized water. The resulting culture plate was incubated at 30 °C for 72 hours at 350 rpm and covered with an adhesive gas permeable membrane (Thermo Fisher Scientific, UK). At the end of the cultivation, the contents of each well was centrifuged. To remove water traces in wells containing beads, the solid phase was recovered and dried under nitrogen gas at room temperature before adding acetone at 1:1 ratio of the cultivation volume and incubating the resulting mixture at 30 °C and 350 rpm for four hours. Following incubation, the mixture was centrifuged and the organic phase recovered for taxadiene analysis via GC-MS. Biomass accumulation was monitored in offline samples as optical density at 600nm using Nanodrop 2000c spectrophotometer (Thermo Fisher Scientific, UK). For comparison, additional cultivations were performed using the widely used dodecane overlay approach described in detail in section 2.7. Control cultivations with YPD only and no extraction aid were also performed, taxadiene was extracted from the solid phase (biomass only) after the cultivation time using acetone at 1:1 ratio of the cultivation volume and incubating the resulting mixture at 30 °C and 350 rpm for four hours. Microscope images were taken using an oil immersion light microscope that is equipped with 100x Leica NPLAN objective lens (Leica Microsystems, Germany) using an Andor-Zyla sCMOS camera (Oxford Instruments, UK). The SPA method using adsorbent HP-20 resin beads was summarized in Figure 2.

**Figure 2.**
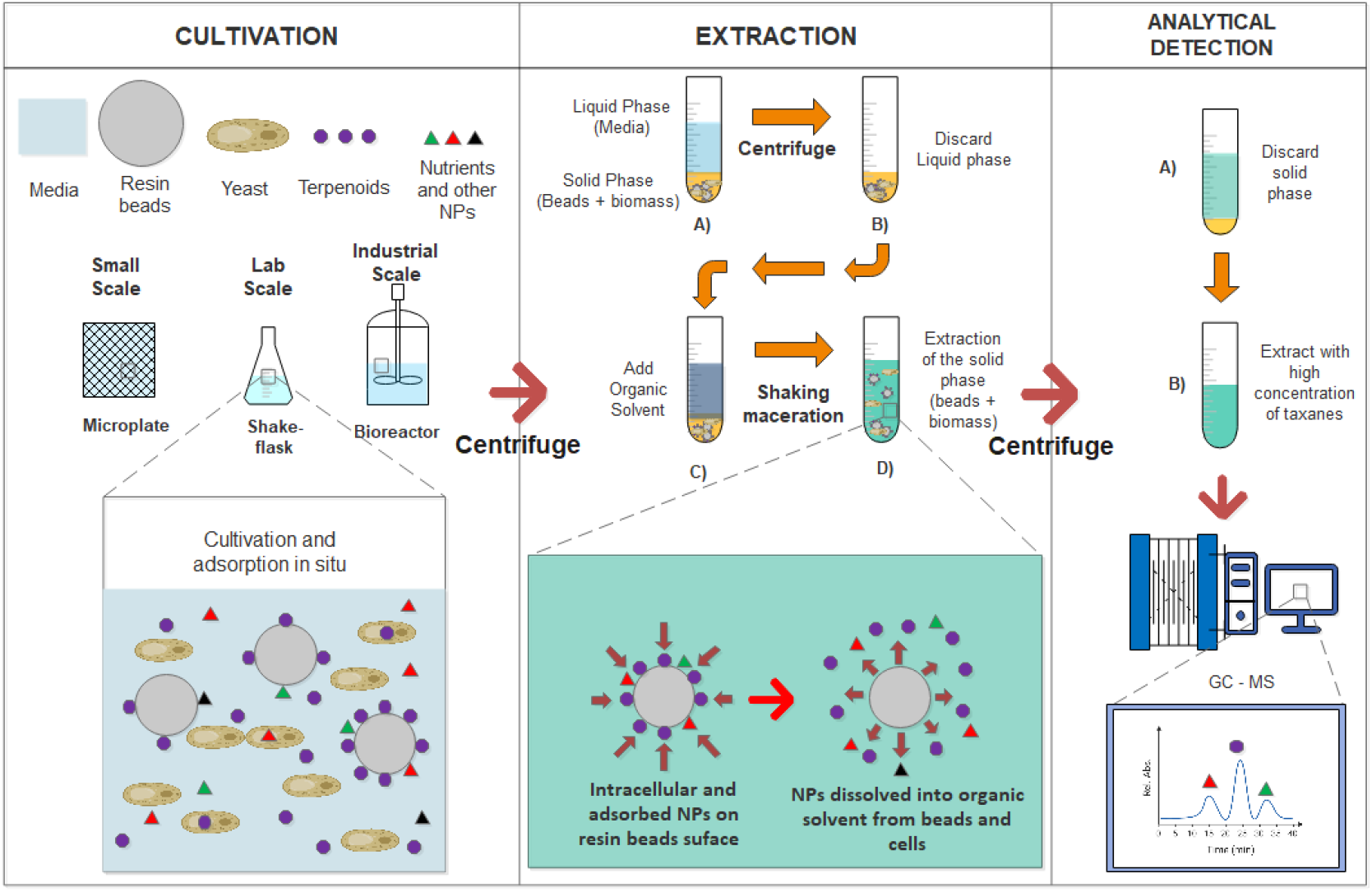
HP-20 resin beads in situ SPA method. The methodology focuses on extracting taxadiene an early paclitaxel precursor produced by the engineered yeast (LRS5). Taxadiene is adsorbed onto the resin surface as it is secreted throughout the cultivation and extracted at the end using an appropriate organic solvent.

### 2.3 Small scale cultivations

Small scale cultivations were performed in 50 mL tubes and 250 mL shake flasks with a working volume of 10 % of their respective total volumes. Inoculum cultures were prepared by transferring a single colony of *LRS5* to 5 mL YPD media and incubating at 30 °C and 250 rpm overnight. An aliquot of this culture was then diluted with YPG media to give a 5 mL culture with an initial OD_600_ = 1. In the SPA cultivations, autoclaved HP-20 adsorbent resin beads were added at a concentration of 3, 6 or 12 % (w/v). All of the cultivations were incubated at 30 °C with a shaking speed of 250 rpm for 72 hours. At the end of the cultivation, biomass accumulation and taxadiene extraction using acetone was performed as described in section 2.2.

### 2.4 Taxadiene partition from solid and liquid phases

To determine the taxadiene partition between the beads, biomass and liquid phase, an additional experiment was conducted under the conditions detailed in Section 2.2. Once the stationary phase of the growth had been reached at around 72 hours the beads were removed from the cultivation using a nylon mesh cell strainer with a pore size of 70 μm (Thermo Fisher Scientific, UK). The resulting solution was then centrifuged and the supernatant transferred to a separate vessel. The biomass and beads were dried under nitrogen gas before being suspended separately in same volume of acetone as they had of cultivation volume and incubated at 30 °C and 250 rpm for four hours. To recover taxadiene from the supernatant, a dodecane overlay of 20 % of the cultivation volume was added to the liquid phase at 30 °C and 250 rpm for four hours. Following incubation, the extracts were centrifuged and the dodecane organic phase was recovered for taxadiene analysis via GC-MS.

### 2.5 Extraction solvent optimization

To investigate the effect of extraction solvent on recovery of taxadiene from the resin beads, the following solvents were tested: ethanol, acetone, ethyl acetate and dodecane were investigated. Following cultivation in small scale culture (5 ml) for 72 hours under the conditions detailed in section 2.3, the solid phase was recovered by centrifugation and re-suspended in a volume of the respective solvent equal to that of the original culture. The resulting mixtures were incubated at 30 °C and 250 rpm for four hours then centrifuged to separate the organic and solid phases. The organic phases from each organic solvent were subsequently extracted for GC-MS analysis.

### 2.6 Bioreactor cultivation

Larger scale cultivations were conducted in MiniBio 500 bioreactors (Applikon Biotechnology, The Netherlands) with a working volume of 250 mL. Pre-inoculum cultures were prepared by transferring from a single colony to 5 mL of YPD and incubating at 30 °C and 250 rpm for eight hours. The resulting culture was subsequently used to inoculate a secondary 10 mL culture to an OD_600_ = 1 and incubated overnight. An aliquot of the resulting culture was diluted with 1 % (v/v) medium to give a 200 mL culture with an initial OD_600_ = 1. To prevent excess foam production polypropylene glycol P2000 (Alfa Aesar, UK) was added to a concentration of 0.01 % (v/v) and a Rushton turbine was placed at the medium-air interface. Autoclaved HP-20 adsorbent resin beads were added at a concentration of 3 % (w/v). A set point of 30 % saturation for DO was applied and the culture temperature was maintained at 30 °C. The pH was maintained above six through the automatic addition of 1M NaOH. Biomass was measured offline twice daily using Nanodrop 2000c spectrophotometer (Thermo Fisher Scientific, UK). Galactose concentration was monitored via offline sampling twice daily using the 3,5-Dinitrosalicylic acid (DNS) method [28].

### 2.7 Dodecane overlay method

Dodecane overlay method was made as point of comparison for our *in situ* SPA method, as it is a methodology widely used to recover taxanes. In this work, the culture, seed culture and extraction were prepared as described before [25] for microscale cultivations. After the cultivation time, an aliquot of dodecane was recovered for taxadiene analysis via GC-MS.

### 2.8 Analytical methods

Taxadiene identification and quantification was achieved via GC-MS. A 1 μL sample of each taxadiene containing organic solvent extract was injected into a TRACE^™^ 1300 Gas Chromatograph (Thermo Fisher Scientific, UK) coupled to an ISQ LT single quadrupole mass spectrometer (Thermo Fisher Scientific, UK). Chromatographic separation was achieved using a Trace Gold TG-SQC gas chromatography column (Thermo Fisher Scientific, UK) using a previously described method [26]. To identify and quantify production of compounds by *LRS5*, pure standards of taxadiene, kindly supplied by the Baran Lab (The Scripps Research Institute, California, USA) and GGOH, obtained from Sigma Aldrich (Gillingham, UK), were used. Characteristic gas chromatograms and taxadiene mass spectra can be found in supplementary information (see supplementary material).

### 2.9 Statistical analysis

Statistical analyses were performed using MATLAB 2020a and Minitab 19 statistical software. One-way analysis of variance (ANOVA) was used to determine whether HP-20 resin beads concentration yielded a significant impact on biomass yield at microplate and shake flask scale. A Dunnett’s multiple comparison test was used to compare each treatment to the control. One-way ANOVA was also used to determine whether the selected extraction solvent had an effect on taxadiene titers recovery. A Tukey’s honest significant difference test was subsequently employed to compare each of the treatments. The null hypothesis considered that there was no significant difference between the treatments, hence if *p* ≤ 0.05 the null hypothesis was rejected.

## 3. Results and discussion

### 3.1 Effect of resin bead concentration on *LRS5* cell growth

*In situ* solid phase adsorption (SPA) using HP-20 resin beads has proven to be an effective method for improving NP recovery in microbial cultivations [16,29]. However, high bead concentrations have been found to inhibit microbial growth [30]. As the rates of biomass and taxane accumulation have been shown to be highly correlated in the engineered *S. cerevisiae* strain, *LRS5*, used in this study [25], optimization was necessary to maximize both growth and product recovery. The *LRS5* strain was cultivated in the presence of a range of bead concentrations in order to determine the effect on growth kinetics at microscale. A further study was conducted using shake flasks to investigate the effect of concentrations at an increased scale. Control cultivations of *LRS5* in the absence of the resin were included in the study, both with or without (no extraction aid) a dodecane overlay. The results of these studies are summarized in Figure 3.

**Figure 3.**
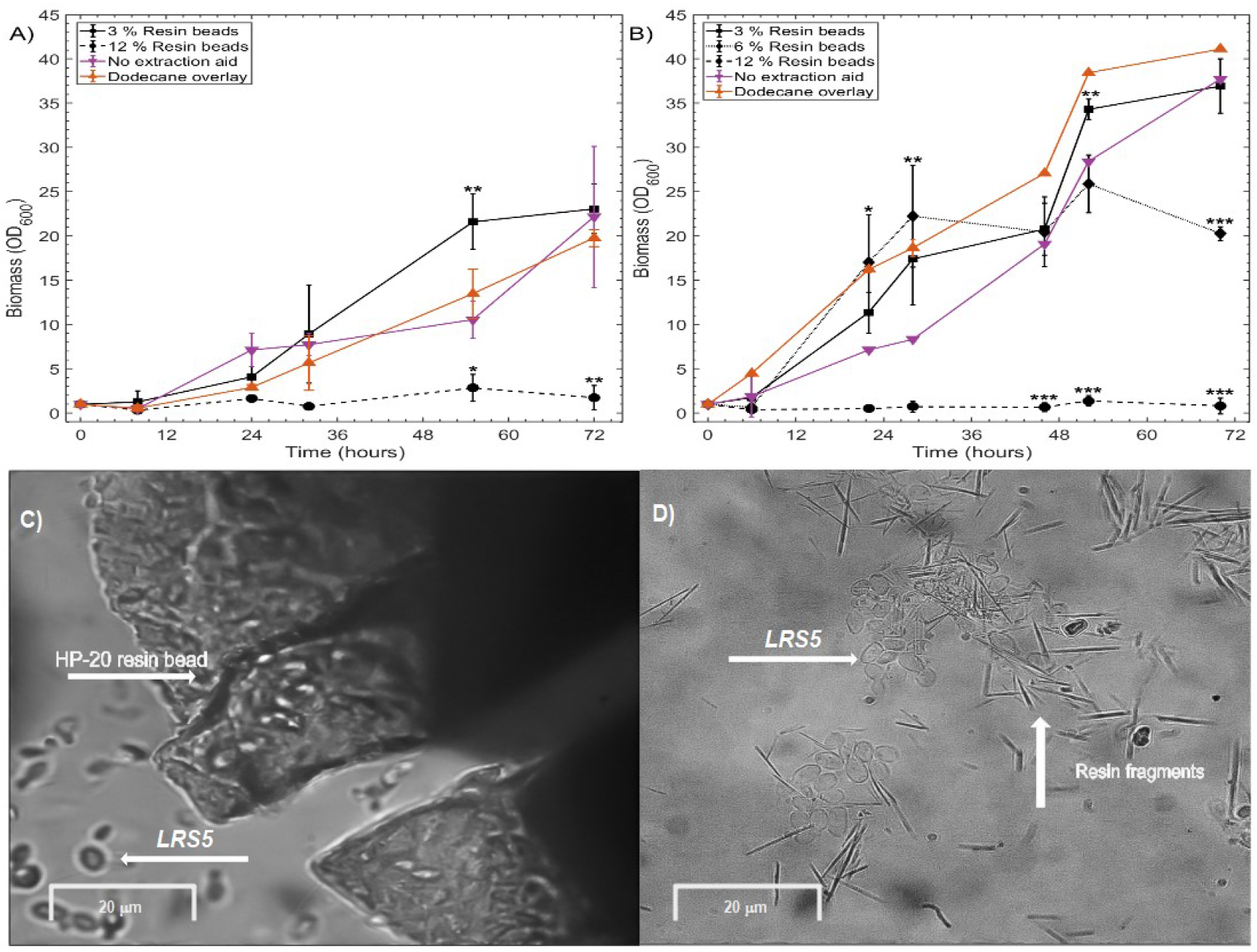
Growth kinetics of yeast strain *LRS5* in presence of different concentrations of HP-20 resin beads (TOP), microscope images of *LRS5* cultivated at 6% (w/v) resin beads at small scale (BOTTOM). 3A) Experiments were carried out using microscale cultivations (2 ml cultivation) and 3B) small scale cultivations (25 ml cultivation). All cultivations were performed with YP media supplemented with 2 % galactose at 30 °C and 250 rpm. Values represent mean ± standard deviation (*n* = 3) (* *p* < 0.05, ** *p* < 0.01, *** *p* < 0.001; Dunnett’s multiple comparison test with no extraction aid (0 % resin bead) as control). 3C) Interaction between HP-20 resin bead and *LRS5* cells. 3D) Sharp resin fragments observed in the culture medium after 72 hours.

At microscale, the addition of 3 % (w/v) HP-20 resin beads was found to enhance the growth rate of *LRS5*, with the stationary phase of growth reached earlier than for the control cultivation with no extraction aid (Figure 3A). An OD_600_ of 21 ± 3 was observed after 55 hours, significantly greater than the control cultivation with no extraction aid (*p* = 0.002). However, the addition of 3 % (w/v) beads had no significant effect on the final OD_600_ (*p* = 0.964) with highly similar OD_600_ values of 23 ± 3 and 22 ± 8 obtained for the 3 % and no extraction aid cultivations. The final biomass yield of the cultures treated with a dodecane overlay was also not statistically different from the control with no extraction aid (*p* = 0.654) at 20 ± 1. At the higher concentration of 12 % (w/v) the addition of beads was highly detrimental to *LRS5* growth and final biomass yield (*p* = 0.004) with an OD_600_ of just 3 ± 1 achieved at the end of the 72 hour cultivations. This indicated that at low concentrations, the beads were not detrimental to *LRS5* growth. To investigate this effect further and determine the optimal concentration of beads a second experiment involving larger shake flask cultivations was performed, here an additional bead concentration of 6 % (w/v) was tested.

As for the microscale cultivations, the final OD_600_ of the 3 % (w/v) bead (*p* = 0.883) cultivations were very similar to the control cultivation with no extraction aid (Figure 3B). At the higher bead concentration of 6 % (w/v) a 46 % reduction in biomass accumulation was observed compared to the control (Figure 3B, *p* < 0.0005). Increasing the concentration further to 12 % (w/v) inhibited growth almost entirely with a final OD_600_ = 1 ± 0.5 (*p* < 0.0005). The optimal bead concentration was therefore deemed to be 3 % (w/v) with maximum OD_600_ values of 21 ± 3 and 37 ± 3 at micro and shake flask scale, respectively.

These results were in agreement with previous studies using *E. coli*, *Streptomyces hygroscopicus* and *Streptomyces virginiae* [16,19,31], which found high bead concentrations were detrimental to microbial growth. *E. coli* was able to tolerate higher bead concentrations with an optimal of 5 % (w/v) [19], whilst the optima for *Streptomyces hygroscopicus* and *Streptomyces virginiae* were similar to that found in this study (Figure 3) at between 2 % and 5 % (w/v) respectively [16,31].

Adsorption to the resin surface is not exclusive to the desired taxadiene compound [32], additional hydrophobic compounds present within the culture medium therefore likely adsorbed to its surface. Such compounds include fatty acids and non-polar amino acids, which are essential for yeast growth [33]. A previous study revealed that 55 % of the casein present within the culture medium adsorbed to the HP-20 beads when a concentration of 20 % (v/v) beads was used compared to just 17 % at a 5 % (v/v) bead concentration [34]. This resulted in a reduced availability of amino acids for cell growth, the adsorption of essential nutrients may have therefore contributed to the poor growth observed in the 12 % (w/v) cultivations of this study. This highlights the importance of carefully optimizing the concentration of resin to ensure sufficient capacity to maximize product recovery whilst minimizing nutrient loss and hence growth inhibition [16]. In addition, visualization of the cultures using oil immersion light microscopy (Figure 3C and 3D) revealed physical degradation of the beads, generating sharp fragments as shown in Figure 3D, which may have caused some mechanical lysis of the cells.

Such degradation was likely enhanced by culture agitation as it has been noted in previous works using similar type of resins [35]. Furthermore, as the resin beads are around 100-fold larger than the yeast cells, their presence within the culture likely contributed to shear stress, which may have hindered cell viability at the higher bead concentrations (>6 % (w/v)). To our knowledge, this is the first report of the contribution of HP-20 bead degradation and subsequent sharp fragment generation to reduced cell growth at higher bead concentrations.

### 3.1 Solid – Liquid Extraction optimization using different organic solvents

The partition of taxadiene between the solid (beads + biomass) and aqueous (culture medium) phases was subsequently investigated in small scale (5 mL) cultures of *LRS5*. Following cultivation, taxadiene was extracted from each phase using dodecane and quantified via GC-MS. Analysis revealed 97 % of the detected taxadiene was present in the solid phase (see supplementary material). As a result, subsequent experiments focused on extraction and purification of taxadiene from the solid phase exclusively. This ensured recovery of the vast majority of the product of interest, whilst minimizing solvent requirements.

A key benefit of the application of an adsorbent resin is the elimination of reliance on biocompatible extraction solvents, which with high boiling points render the purification stage both energy intensive and costly (*i.e*. dodecane). A low cost extraction solvent with a low boiling point is preferable to maximize the environmental and economic sustainability of the process. Previous studies have shown solvents with intermediate polarity such as acetone can improve terpenoid recovery in *in situ* SPA systems compared to dodecane [36,37]. Aguilar et al. (2019), recently employed HP-20 resin beads for the *in situ* extraction of the sesquiterpene zizaene from engineered *E. coli* cultivation. In that study, of the seven extraction solvents tested in the study, ethyl acetate, decane and isooctane displayed superior zizaene recovery to dodecane. Ethyl acetate has also been successfully employed for the recovery of limonene [38] and Fusicocca-2,10(14)-Diene [39] from resins resulting from *E. coli* and *S. cerevisiae* cultivations, respectively. Ethyl acetate was therefore selected as a potential solvent in this study, as in addition to its observed affinity for terpene compounds, it is a volatile and relatively low-cost solvent with low toxicity. Acetone and ethanol with similar and higher polarities compared to ethyl acetate, respectively, are commonly used organic solvents for the extraction of carotenoids from yeast cells [40]. The efficacy of these solvents for eluting the shorter chain diterpene, taxadiene, from the HP-20 resin was investigated in this study, as summarized in Figure 4.

**Figure 4.**
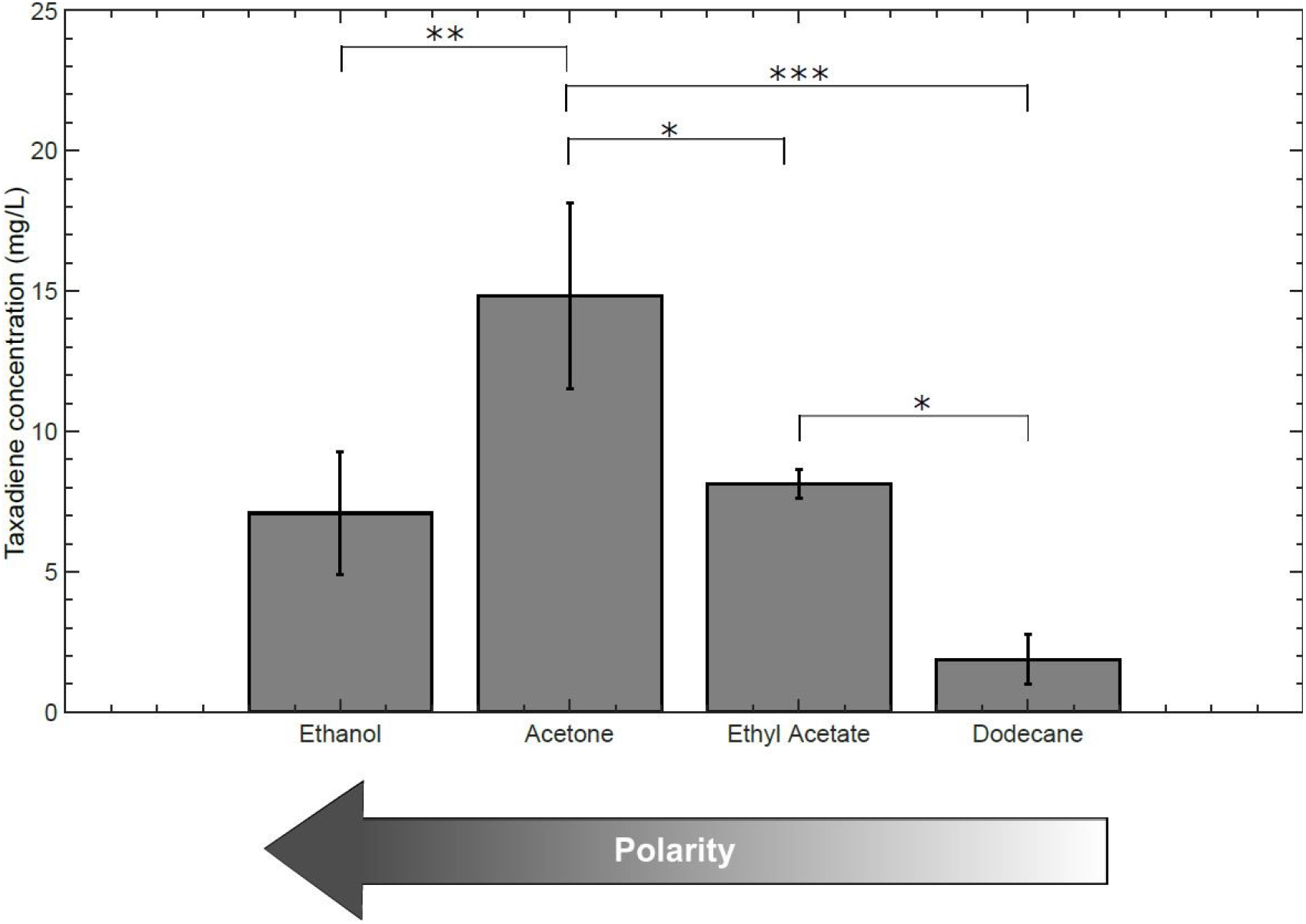
Effect of different organic solvents on taxadiene recovery in Solid phase adsorption (SPA) cultivation method. All the solvents were used in the extraction stage, following resin bead cultivation at small scale (5 ml). Values represent mean ± standard deviation (*n* = 3) (* *p* < 0.05, ** *p* < 0.01, *p* < 0.001; Tukey’s honestly significant difference multiple comparison test).

Of the tested solvents acetone (*p* = 0.007) and ethyl acetate (*p* = 0.023) enhanced taxadiene recovery significantly with respect to dodecane as shown in Figure 4. Unlike dodecane, the three other solvents tested in this investigation are not biocompatible and are known to affect yeast-cell viability and metabolic activity retention at concentrations from 6 % (v/v) [41]. Organic solvents including ethanol, acetone and ethyl acetate, penetrate the lipid portion of the cell membrane, which causes permeabilization and can ultimately lead to cell lysis [42–44]. Biocompatible solvents such as dodecane are commonly used for *in situ* liquid-liquid extraction to minimize this effect during cultivation and maximize cell viability. However, permeabilization is desirable for maximizing product recovery as it ensures access to intracellular taxadiene [42,45]. The higher titers observed with the alternative solvents in this study (Figure 4) was likely attributed to increased membrane permeability and intracellular taxadiene recovery. The greatest taxadiene titer of 15 ± 2.0 mg/L was achieved using acetone (5 ml cultivations), four-fold greater than that using the highly non-polar and biocompatible dodecane solvent. This solvent was therefore selected for subsequent studies.

### 3.2 Synergic effect of HP-20 resin beads and selected solvent to improve taxadiene extraction

The results of the desorption step indicated that the use of ethanol, acetone or ethyl acetate as the extraction solvent, resulted in higher taxadiene titres than dodecane. Such solvents are known to increase cell membrane permeability [43,44] and previous studies have shown that some of the taxadiene is retained intracellularly [25]. It was therefore hypothosised that improved recovery achived using ethanol, acetone or ethyl acetate may be partially due to increased cell memebrane permeability. In order to investigate this further, an additional experiment was performed using the highest yielding solvent, acetone (Figure 4). *LRS5* was cultivated in the absence of resin beads or a dodecane overlay (no extraction aid) and acetone was applied at the end of the cultivation to extract taxadiene from the solid phase (biomass). This was compared to the optimal *in situ* solid phase (beads + biomass) cultivation with 3 % (w/v) HP-20 beads and the traditional dodecane overlay method as shown in Figure 5.

**Figure 5.**
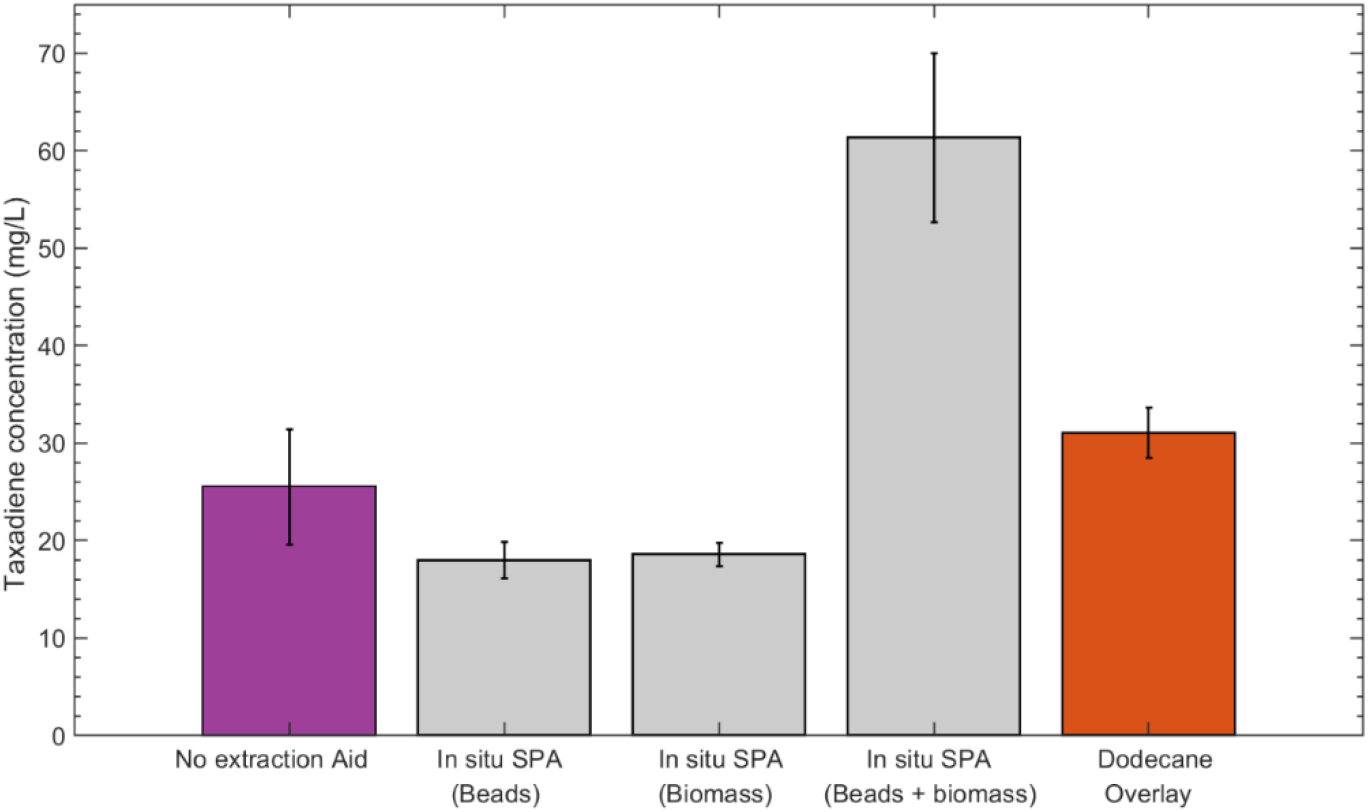
Taxadiene recovery titers (bars) from microplate cultivations (2 mL cultivation volume) after 72 hours of cultivation at 30 °C and 350 rpm. Purple bar represents the cultivation made with no extraction aid where the extraction was made to the biomass using acetone. Colored grey bars represent *in situ* Solid phase adsorption (SPA) cultivation using 3 % (w/v) bead concentration with further extractions made with acetone whilst orange bar represents extraction made with dodecane using the dodecane overlay method. *LRS5* reached similar OD at 600 nm in all treatments after 72 hours. Beads + Biomass value could be higher due to less manipulation in the extraction steps. Values represent mean ± standard deviation (*n* = 2).

Biomass accumulation was similar for the SPA, no extraction aid and dodecane overlay cultivations with final OD_600_ values of 23 ± 1, 22 ± 3 and 20 ± 1, respectively. When acetone was applied to extract taxadiene from the biomass resulting from the control treatment, in which *LRS5* was grown in the absence of an *in situ* extraction aid, the final taxadiene titer was 26 ± 6 mg/L at microscale as shown in Figure 5. Interestingly, this was comparable to the 31 ± 3 mg/L obtained using the traditional dodecane liquid-liquid extraction method. This suggested that, in the absence of an *in situ* extraction aid, a significant proportion of the taxadiene was intracellular or adsorbed onto cell surface rather than secreted, likely due to the hydrophobic nature of the diterpene. In the *in situ* SPA cultivations, taxadiene was distributed almost equally between the resin beads and biomass with titers of 18 ± 2 mg/L and 19 ± 1 mg/L observed, respectively. This indicated that the cells retained up to 50 % of the taxadiene produced during the cultivation. However, it was found that during the separation and desorption steps, around 30% of the bead volume was lost compared to parallel cultivations with the same bead concentration, in which the beads and biomass were not treated separately (see supplementary material). The 50 % taxadiene repartition between biomass and resin beads provides an important piece of information, as it showcases where the taxadiene that is being detected in the system is being captured. To our knowledge, this information has not been commonly reported as the extraction is generally made to the biomass and resin in conjunction [16,29,46] or to the single beads without biomass [17,47]. These results could have an important impact into determining the capacity of the resin beads to relieve the cells from intracellular toxicity of selected NP among other phenomena.

Simultaneous extraction of taxadiene from the resin beads and biomass, without separation, resulted in a taxadiene titer of 61 ± 8 mg/L, at this scale. This suggests that around 39 % of the recoverable taxadiene was lost during the separation step, likely due to the loss of beads during the biomass and beads separation, indicating that the actual fraction of taxadiene retained by the cells was likely around 50 %. This is in agreement with a previous study using *LRS5*, which demonstrated that up to one third of the total taxadiene was retained intracellularly using dodecane [25]. Using the optimal bead concentration (3 % (w/v)) and extraction solvent (acetone) resulted in a 1.8-fold improvement in taxadiene recovery compared to the widely used dodecane overlay approach (Figure 5). Therefore, to maximize intracellular and secreted taxadiene titer, an extraction solvent with a high affinity for taxadiene and a high permeabilization ability should be selected following the cultivation. Differences in results between Figure 4 and Figure 5 were likely the result of an increased shaking speed of 350 rpm (Figure 5) compared to the 250 rpm applied for the 5 mL cultures (Figure 4). In addition, the microtiter plates were sealed with a gas permeable membrane whereas the tubes used for the 5 mL cultures were airtight. Oxygen mass transfer was therefore likely enhanced significantly at microscale, promoting aerobic growth and thus improving taxadiene titers. A similar effect was observed in a previous study where taxadiene titers were improved two-fold in baffled microtiter plate compared to shake flask cultivations of the strain [21].

A significant quantity of taxadiene was recovered from the cells resulting from the cultivation in the absence of an extraction aid using acetone (Figure 5). This indicated that, in addition to promoting elution of secreted taxadiene from the HP-20 resin, acetone also enhanced the recovery of intracellular taxadiene, likely due to its effect on cellular permeability. The proposed SPA method is therefore an integrated bioprocess, eliminating the requirement for an additional unit operation for cellular lysis [1,48] or further metabolic engineering to introduce efflux pumps (e.g. outer membrane protein TolC) [49] to enhance the product secretion.

### 3.3 Scale up bioprocess of the *in situ* SPA method

In order to validate the proposed *in situ* SPA method under industrially relevant conditions, the process was scaled up using MiniBio 500 bioreactors (Applikon Biotechnology, The Netherlands). Here industrially relevant process parameters including pH, temperature and dissolved oxygen concentration were monitored and controlled online in real-time, allowing industrial scale conditions to be more effectively mimicked [50]. The results of this experiment are summarized in Figure 6.

**Figure 6.**
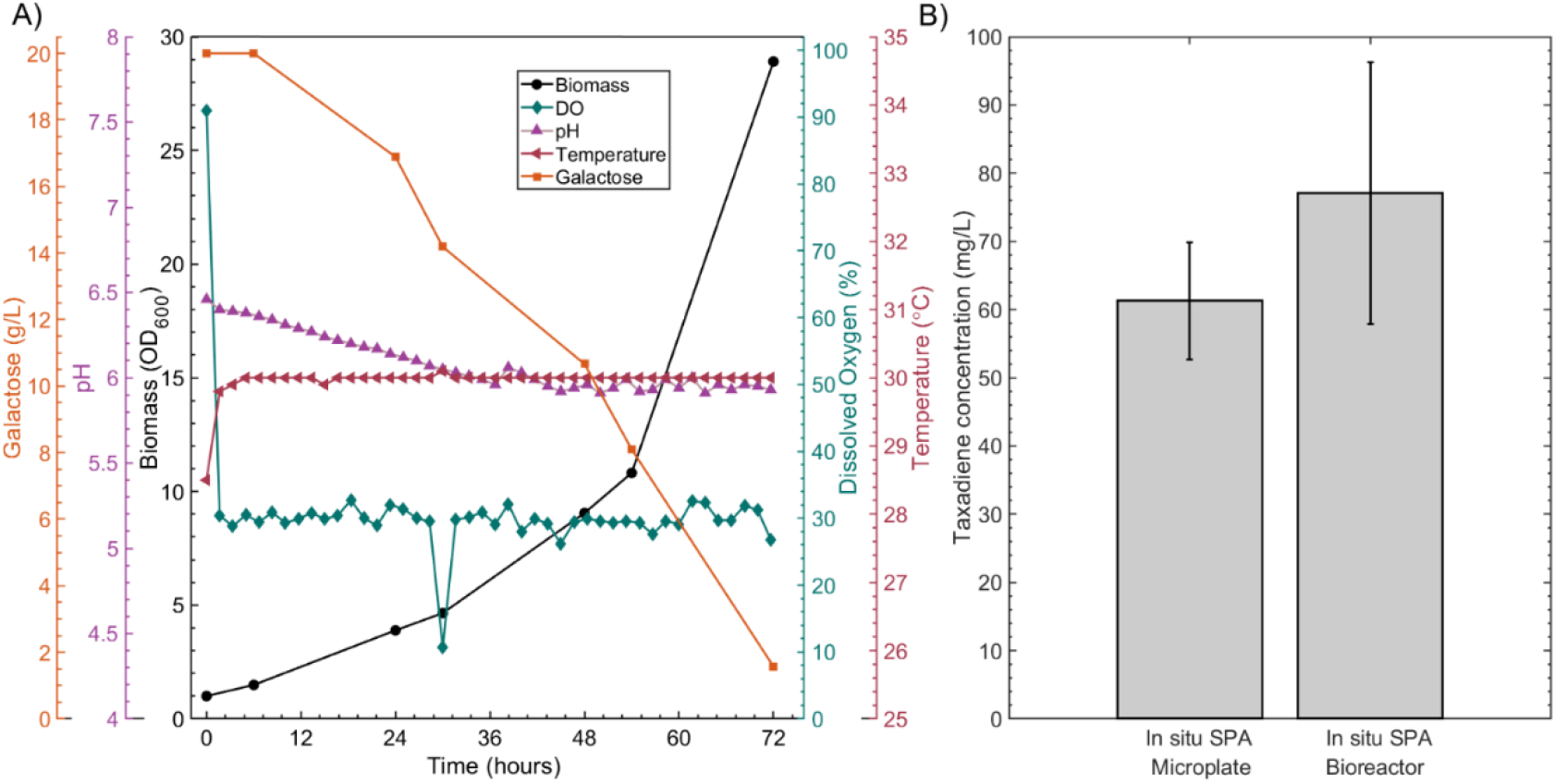
Bioreactor cultivation kinetic parameters of *LRS5* using the *in situ* Solid phase adsorption (SPA) cultivation method. Figure 6A) shows the cultivation parameters using the *in situ* SPA method in YPD media for 250 ml cultivation volume. The controlled parameters were; pH, temperature and dissolved oxygen. Figure 6B) shows the taxadiene titers for microplate cultivation and bioreactor using the *in situ* SPA method using 3 % (w/v) and acetone in the extraction step. All cultivations conditions were the same in these experiments, with the difference that in the bioreactor cultivation pH, temperature and dissolved oxygen parameters were controlled by the equipment. Values in figure 6B represent mean ± standard deviation (*n* = 2).

Taxadiene titers were enhanced at bioreactor scale compared to previous microscale (Figure 6) cultivations using the *in situ* SPA method. Here, the oscillations in key process parameters such as pH and dissolved oxygen could be minimized through automated feedback control facilitating improved taxadiene production. A taxadiene titer of 76 ± 19 mg/L was achieved, compared to 61 mg/L at microscale (Figure 6). This also represented a 1.4-fold improvement in titer compared to liquid-liquid extraction using a dodecane overlay under similar cultivation conditions [25]. A taxadiene titer of 53 mg/L was observed for the *LRS5* strain cultivated in the bioreactor with a dodecane overlay at 30 °C [25]. One possible reason for the enhanced recovery is the positive effect of acetone for solid phase (biomass + beads) extraction, allowing recovery of both the intracellular and secreted taxadiene. Moreover, the loss of taxadiene due to air stripping or co-evaporation with dodecane was avoided through the use of HP-20 resin beads.

These results demonstrate the potential of the *in situ* SPA cultivation method for effective recovery of taxadiene under industrially relevant conditions with improved sustainability.

## 4. Conclusions

The application of this novel method at micro and bioreactor scale improved taxadiene recovery 1.4 and 1.9-fold, respectively, compared to the traditional liquid-liquid extraction approach. Higher resin bead concentrations (from 12 % w/v) resulted in bead fragmentation (mechanical cell disruption) which were found to be detrimental to cellular growth. Acetone ensured the sequestration of both the intracellular and secreted taxadiene at the extraction step achieving process integration. In addition, it was found that from the total taxadiene that is being synthesized, 50 % is adsorbed by the resin beads and the rest is retained in the biomass.

## Acknowledgements

Authors would like to thank the technicians who helped with the GC-MS analyses MRes Stuart Martin and MSc Caroline Delahoyde. Thanks for the support from the IBioE technician Miss Katalin Kis. Thanks to Professor Phil Baran’s Lab at The Scripps Research Institute, San Diego, California for providing the taxadiene standard.

