## Supplemental Material for "*In situ* solid-liquid extraction enhances recovery of taxadiene from engineered *S. cerevisiae* cell factories"

### Supplementary Material

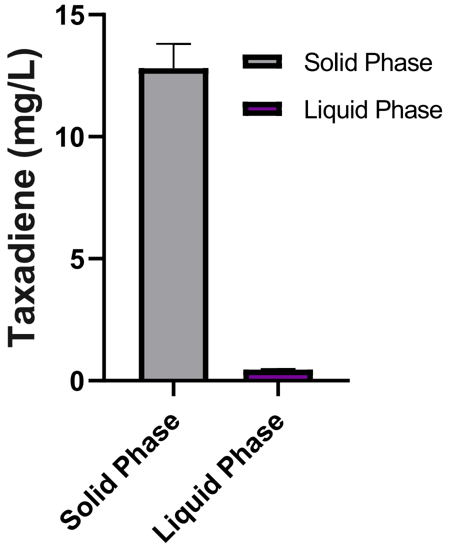


Figure S1. Taxadiene repartition in the culture. Cells, beads and liquid (media) phase Taxadiene concentration comparison after the cultivation. In situ solid phase adsorption method cultivation method in falcon tube (5 mL) was used at 30 ºC. Solid and liquid phase were treated with dodecane in the extraction (see section 2.4 for more details).

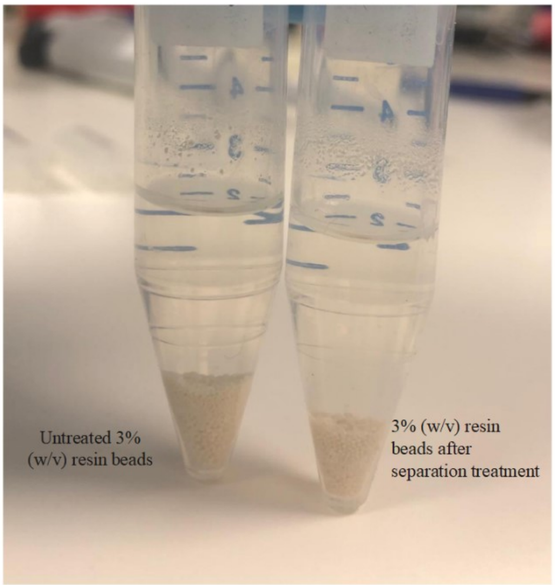


Figure S2. Falcon tubes showing resin beads concentration. On left, the tube shows more quantity of beads as it has not been manipulated, this represents resin beads concentration 3% (w/v) of the microplate cultures. On right it is the quantity of resin beads after the separation of the cells and beads, this represents a considerable lost for around 30%.

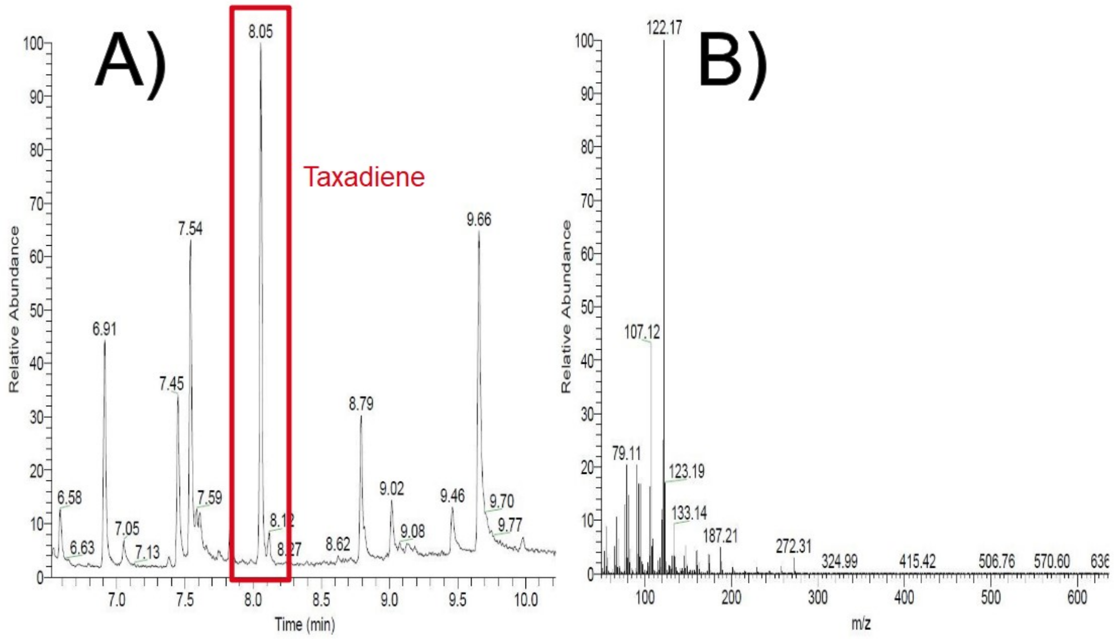


Figure S3. GC-MS Chromatogram A) and Mass spectra B) for microscale cultivation 2 mL using in situ solid phase adsorption method and acetone. Taxadiene peak can be seen at a retention time between 8.05 and 8.08 minutes (figure S2A). Mass spectra is showing the ionization composition of taxadiene. This configuration was maintained for all cultivations scales.

Table S1. Loading plan of the design of experiment for microplate in situ solid phase adsorption cultivation of LRS5 strains.

| **Well** | **Yeast strain** | **HP20 resin bead (% (v/v))** | **Extraction time (hours after fermentation)** | **Dodecane volume (mL)** |
|---|---|---|---|---|
| A1 | *LRS5* | 20 | 24 | 0 |
| A2 | *LRS5* | 5 | 3 | 0 |
| A3 | *LRS5* | 5 | 24 | 0 |
| A4 | *LRS5* | 20 | 3 | 0 |
| A5 | *LRS5* | 5 | 3 | 0 |
| A6 | *LRS5* | 20 | 3 | 0 |
| B1 | *LRS5* | 5 | 24 | 0 |
| B2 | *LRS5* | 20 | 24 | 0 |
| B3 | *LRS5* | 20 | 3 | 0 |
| B4 | *LRS5* | 5 | 3 | 0 |
| B5 | *LRS5* | 5 | 24 | 0 |
| B6 | *LRS5* | 20 | 24 | 0 |
| C1 | *LRS5* | - | 3 | 0 |
| C2 | *LRS5* | - | 3 | 0 |
| C3 | *LRS5* | - | 3 | 0 |
| C4 | *LRS5* | - | 0 | 0.6 |
| C5 | *LRS5* | - | 0 | 0.6 |
| C6 | *LRS5* | - | 0 | 0.6 |
| D1 | None | 5 | 3 | 0 |
| D2 | None (control) | 5 | 3 | 0 |
| D3 | None (control) | 5 | 3 | 0 |
| D4 | None (control) | - | - | 0 |
| D5 | None (control) | - | - | 0 |
| D6 | None (control) | - | - | 0 |
